# Investigating the factors influencing antibiotic use practices and their association with antimicrobial resistance awareness among poultry farmers in Enugu State, Nigeria

**DOI:** 10.1101/2025.02.08.637249

**Authors:** Chika P. Ejikeugwu, Emmanuel A. Nwakaeze, Chikaodi W. Aniekwe, Euslar N. Onu, Michael U. Adikwu, Peter M. Eze

## Abstract

**Background:** The irrational use of antibiotics in poultry production has far-reaching consequences and continues to impact the fight against antimicrobial resistance (AMR) in Africa. In Nigeria, antibiotics are available over-the-counter and are widely used in food animal production for various reasons, including prophylaxis and growth promotion. While this practice may support animal production, it also drives the spread of AMR, posing serious health challenges due to close human-livestock interactions and the country’s high disease burden. This study examined poultry farmers’ knowledge, attitudes, and practices (KAP) regarding antibiotic use and AMR, aiming to highlight the public health risks and challenges in combating AMR in Nigeria.

**Methods:** A cross-sectional survey of 200 poultry farms in Enugu State, Southeast Nigeria, was conducted to evaluate farmers’ KAP towards antibiotic usage and AMR. Using a validated, standardized, and self-administered questionnaire, data were collected from farmers responsible for key farm decisions, including input and feed management. The questionnaire comprised three sections: socio-demographic data, knowledge of AMR, and knowledge and practices regarding antibiotic use. Ethical approval was obtained, and participants gave oral consent based on their professional capacity.

**Results:** The evaluation of poultry’s farmers KAP regarding antibiotic usage and AMR revealed that the majority of farmers (90.5%) reported using antibiotics, primarily for treating infections (80.5%) or for feed enhancement, growth promotion, and prophylaxis (61%). Ampicillin (75%), ciprofloxacin (71.5%), and doxycycline (71%) were the most commonly administered antibiotics. Monthly administration was the most common (48%), and 89% of respondents believed that antibiotics promote poultry growth. Interestingly, a substantial proportion of respondents (65%) were unaware of AMR, highlighting a significant knowledge gap, with only 16% recognizing the risk of AMR infections.

**Conclusion:** Our study revealed that the surveyed poultry farmers heavily rely on antibiotics, primarily for treating infections and, to a significant extent, for growth promotion, despite their limited awareness of AMR. Ampicillin was identified as the most commonly used antibiotic, raising concerns due to its potential link to beta-lactamase selection amid the country’s carbapenem resistance issues. These findings underscore the urgent need for targeted education to address the AMR knowledge gap and reduce the misuse of antibiotics in poultry farming settings in Nigeria.

## INTRODUCTION

Antimicrobial resistance (AMR) is a rapidly evolving and significant global public health threat, contributing to numerous deaths worldwide. The misuse and overuse of antibiotics, particularly in animal production systems such as poultry farming, are key factors driving the persistence of AMR, especially multidrug-resistant phenotypes, such as those producing carbapenemases (e.g. metallo-beta-lactamases), in the Nigerian environment [1,2]. Reducing or eliminating the use of antibiotics in animal production could play a crucial role in mitigating the spread of AMR. To achieve effective interventions, it is essential to address critical ecological gaps and improve knowledge of antibiotic use practices and AMR, particularly within the food production sector. Numerous studies have highlighted the role of antibiotic use in food animal production as a primary driver of the emergence and dissemination of AMR, resistant bacteria and antibiotic resistance genes (ARGs) in the environment [1-4].

AMR presents a growing concern for both human and animal health, as the overuse of antibiotics accelerates the development of resistance in bacterial populations. Approximately 75% of antibiotics sold globally are used in livestock production, often as growth promoters or for disease prevention and treatment [4-6]. Current projections estimate that by 2050, AMR could be responsible for 10 million deaths annually [5]. Given the increasing prevalence of AMR, particularly in developing countries [1,7], it is imperative to understand the factors contributing to the widespread and often unregulated use of antibiotics in agricultural systems, such as poultry farming in Nigeria.

The use of antibiotics in animal production is a constant source of antibiotics that pollutes the environment particularly the soil – where the soil resistome may be impacted to perpetuate the evolution and spread of antibiotic-resistant bacteria and genes [6-9]. In Nigeria, as in many developing countries in Africa, poultry farms may serve as a significant anthropogenic source of environmental antibiotic contamination, as antibiotics are frequently sold over the counter without prescription [1,10]. Many poultry farmers, who typically lack formal training in poultry farming, are at risk of misusing antibiotics, inadvertently contributing to the rising prevalence of AMR in the environment [11-13]. Often, antibiotic use in poultry farming is not based on veterinary recommendations, and non-compliance with withdrawal periods prior to slaughter increases the likelihood of antibiotic residues, ARGs, and resistant bacteria entering the food supply. Poultry farms, therefore, serve as important reservoirs for the development and spread of AMR and resistance genes [1,10].

Although studies have investigated the prevalence of AMR and ARGs in poultry production environments in Nigeria [14], there is limited research on poultry farmers’ perspectives regarding antibiotic use and AMR, particularly in Enugu State, Southeast Nigeria. The use of antibiotics in poultry production also leads to environmental contamination when manure from antibiotic-fed animals is used as fertilizer, contributing to the abundance and spread of resistant bacteria and ARGs through runoff and leaching into water systems [6-9]. This highlights the need for a deeper understanding of farmers’ knowledge, attitudes, and practices (KAP) regarding antibiotic use and AMR in Southeast Nigeria to inform strategies aimed at reducing antibiotic use and preventing the spread of resistance in the region.

As poultry farming is a vital part of Nigeria’s agricultural economy, assessing poultry farmers’ knowledge, behavior, and attitudes toward antibiotic use and AMR is essential to safeguarding antibiotic efficacy, protecting public health, and mitigating the spread of resistance. Previous studies have shown that farmer knowledge and behavior significantly influence antibiotic use decisions [10]. Addressing these factors through targeted training programs and interventions could promote more responsible antibiotic use, ultimately reducing AMR spread in poultry production systems [11,12].

The goal of this study was to investigate the drivers of irrational antibiotic use in poultry farming, which contribute to the proliferation of AMR in the environment. A validated, standardized, and structured questionnaire comprising both open-ended and closed-ended questions was employed to collect critical data from poultry farmers, including socio-demographic information, knowledge of AMR, and practices regarding antibiotic use. To minimize bias, face-to-face interviews were conducted with individuals responsible for farm management decisions. The findings of this research are expected to bridge existing knowledge gaps by assessing the KAP of poultry farmers regarding antibiotic use and AMR in Enugu State, Southeast Nigeria, and to highlight and contribute to mitigating the public health risks posed by inadequate awareness of AMR and irrational antibiotic application in poultry.

## MATERIALS AND METHODS

### Study design

The study was conducted from April 1 to August 31, 2024. A cross-sectional design was used, and 200 privately owned poultry farms (each with at least 100 birds) were randomly selected for study. The study assessed the poultry farmers responsible for key agricultural decisions, such as input usage, feed management, and procurement.

### Study location

A total of 200 privately owned farms were randomly selected from nine different locations across three senatorial zones of Enugu State: Enugu East, Enugu West, and Enugu North (Figure 1). These senatorial zones in Enugu State are renowned for their prominence in livestock farming, particularly poultry.

**Figure 1.**
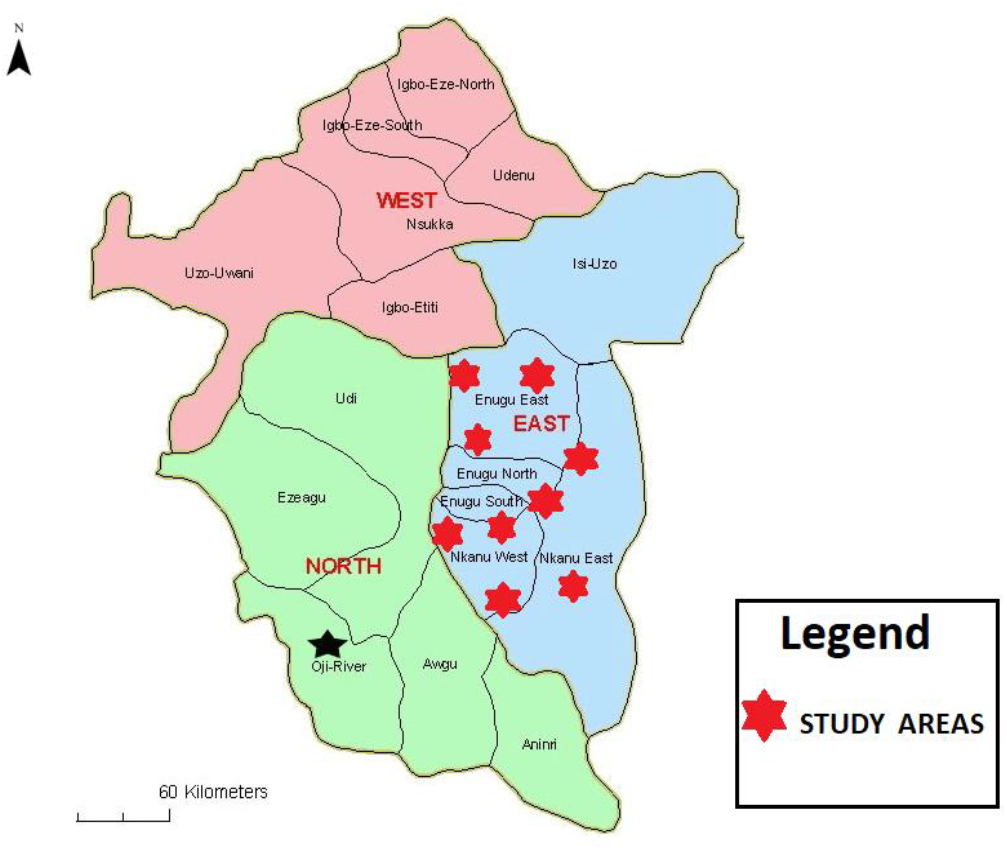
Map of the studied location in Enugu State, Nigeria (Adapted from Enugu Connect [15])

### Study Instrument

The study instrument was a validated, standardized, and self-administered questionnaire. It was available in English and designed to collect essential, farmer-specific data, including socio-demographic information, knowledge of AMR, and practices regarding antibiotic use. Guided by existing literature on farmer KAP towards AMR and antibiotic use [12-14], the questionnaire was structured into three main sections: (1) Socio-demographic data, including age, gender, and educational background; (2) Knowledge of AMR; and (3) Knowledge and practices regarding antibiotic use. The questionnaire underwent a pre-testing phase, conducted with the assistance of a veterinarian to ensure clarity, relevance, and accuracy of content. Necessary revisions were made based on feedback from the pre-test, improving the tool’s effectiveness for subsequent data collection across the surveyed farms. All data of the respondents (poultry farmers) in this study remained anonymous to the researchers/authors. The responses were collected in categorical (yes, no, no idea or no response), ordinal (e.g., agree, disagree, strongly agree, strongly disagree), and open-ended formats to allow for both quantitative and qualitative analysis. Subsequently, the term “respondents” shall be used interchangeably with “poultry farmers” in this study. All responses from the interviews were documented on-site.

### Data Management and Analysis

Data were collated and analyzed using Microsoft Excel version 16.92 (24120731). Categorical variables were summarized as frequencies and percentages to provide insights into prevalent farming practices, as well as the respondents’ knowledge and awareness levels concerning AMR and antibiotic use. A two-way ANOVA was performed to evaluate statistical significance across the respondent outcomes.

### Ethical approval

Ethical approval for the study was granted by the Enugu State Ministry of Health (Reference number: MH/MSD/REC21/630). Informed oral consent was obtained from all study participants. Additionally, participants provided consent for the publication of the study data.

## RESULTS

### Socio-demographic features of poultry farmers

This study included 200 private-owned poultry farms (with at least <100 birds) in Enugu, Southeast Nigeria. Table 1 summarizes the socio-demographic characteristics of the study population, covering aspects such as gender, age, marital status, occupation, education, and internet usage, all of which may influence the KAP related to antibiotic use and AMR among the surveyed farmers. This socio-demographic overview underscores the diversity and digital connectivity of the studied population, both of which are crucial in understanding their attitudes toward antibiotic use and AMR.

**Table 1:** Distribution of the studied respondents according to socio-demographic data.

| Demographics | Variable | n | % |
| --- | --- | --- | --- |
| Gender | Male | 88 | 44 |
|  | Female | 112 | 56 |
|  | **Total** | 200 | 100 |
| Age | <20 years | 8 | 4 |
|  | 21 – 30 years | 54 | 27 |
|  | 31 – 40 years | 50 | 25 |
|  | 41 – 50 years | 44 | 22 |
|  | 51-60 years | 21 | 10.5 |
|  | >60 years | 23 | 11.5 |
|  | **Total** | 200 | 100 |
| Marital status | Single | 66 | 33 |
|  | Married | 132 | 66 |
|  | Divorced | 1 | 0.5 |
|  | No response | 1 | 0.5 |
|  | **Total** | 200 | 100 |
| Occupational status | Employed | 74 | 37 |
|  | Self Employed | 42 | 21 |
|  | Housewife | 16 | 8 |
|  | Househusband | 1 | 0.5 |
|  | Student | 36 | 18 |
|  | Retired | 16 | 8 |
|  | Others | 14 | 7 |
|  | No response | 1 | 0.5 |
|  | **Total** | 200 | 100 |
| Highest academic qualification | Primary | 5 | 2.5 |
|  | Secondary School | 62 | 31 |
|  | University | 76 | 38 |
|  | Diploma | 12 | 6 |
|  | OND/HND | 11 | 5.5 |
|  | Masters | 18 | 9 |
|  | Doctorate | 8 | 4 |
|  | No response | 8 | 4 |
|  | **Total** | 200 | 100 |
| Internet usage | Everyday | 108 | 54 |
|  | Often | 57 | 28.5 |
|  | Rarely | 21 | 10.5 |
|  | Never | 13 | 6.5 |
|  | No response | 1 | 0.5 |
|  | **Total** | 200 | 100 |

There was a slightly higher proportion of female respondents (56%) compared to male (44%), achieving a balanced gender representation essential for capturing diverse perspectives on AMR and antibiotic use. Age distribution reveals that most respondents (52%) are between 21 and 40 years, suggesting the study primarily reflects the viewpoints of younger to middle-aged adults who are mainly involved in poultry production business in Enugu, Nigeria. Representation is lower for those under 20 years (4%) and over 60 years (11.5%) indicating lesser participation from these age groups (Table 1).

In assessing the relationship between poultry farmers’ knowledge of antibiotic use and their socio-demographic characteristics, we found no significant differences across various groups for all variables (p>0.05; Table S1). Among male respondents, 93.18% demonstrated knowledge about antibiotic use, while 97.32% of female respondents showed similar knowledge. The p-value of 0.570 indicates that gender does not significantly impact knowledge levels (p > 0.05). Knowledge of antibiotic use was consistently high across all age groups, ranging from 90.91% among respondents aged 41-50 years to 100% in several other age groups, with a p-value of 0.392, showing no significant age-related differences (p > 0.05; Table S1).

In terms of marital status, a significant majority of respondents are married (66%), followed by single individuals (33%). Minimal representation is observed among divorced respondents and non-respondents (0.5% each). Marital status was also not a significant factor since 98.48% of married respondents and 90.91% of single respondents displayed knowledge about antibiotic use, with a p-value of 0.487 (p > 0.05). Similarly, occupational status did not significantly influence knowledge levels, as knowledge ranged from 91.67% among students to 100% among housewives and househusbands, with a p-value of 0.448 (p > 0.05).

The occupational status is varied, with most participants being employed (37%) or self-employed (21%). Students make up 18% of the sample, while housewives and retired individuals each account for 8%. Educational attainment showed no significant association with knowledge of antibiotic use: knowledge levels ranged from 80% among primary school graduates to 100% among respondents with OND/HND and Doctorate degrees (p = 0.457; p > 0.05). Education levels of the respondents are relatively high, with 38% holding a university degree. The presence of respondents with only primary education (2.5%) is minimal, suggesting that the respondents are relatively well-educated overall (Table 1).

Internet usage amongst the respondents is frequent, with 54% accessing it daily and 28.5% often, indicating high digital engagement. Furthermore, internet usage frequency did not affect knowledge significantly; 90.48% of respondents who rarely used the internet and 98.25% of those who used it frequently were knowledgeable about antibiotic use, with a p-value of 0.467 (p > 0.05). Overall, knowledge levels were high across all socio-demographic variables, suggesting a broadly informed group of poultry farmers (Table S1).

### KAP of poultry farmers regarding antibiotic application in poultry farms

When asked if they know the class, names or brands of the antibiotics administered to their birds, about 80% of the respondents responded in affirmation. While a small proportion of respondents (2.5%) applied antibiotics at all times, a significant number of the respondents applied antibiotics to their birds when there are no symptoms (24%) or when there are signs of infections (77%). Only 15% of the respondents follow the recommendations of a veterinarian in the administration of antibiotics to their poultry birds (Table S2).

When asked about the frequency of application and most used antibiotics, 48% of the respondents applied antibiotics on a monthly basis while 29.5% respondents affirmed that they applied antibiotics bi-weekly. Ampicillin (75%) was notable as the most applied antibiotics, followed by ciprofloxacin (71.5%), gentamicin (67%) and amoxicillin (59%).

When asked about the reasons for administering antibiotics a significant proportion of the respondents (80.5%) used antibiotics to treat infections rather than for growth promotion (12.5%). Only 43.5% of the respondents applied antibiotics to their birds for prophylactic measures. Surprisingly, a significant proportion of the respondents affirmed that antibiotics are good for the promotion of growth in poultry birds (89%) (Figure 2 and 3; Table S2).

**Figure 2.**
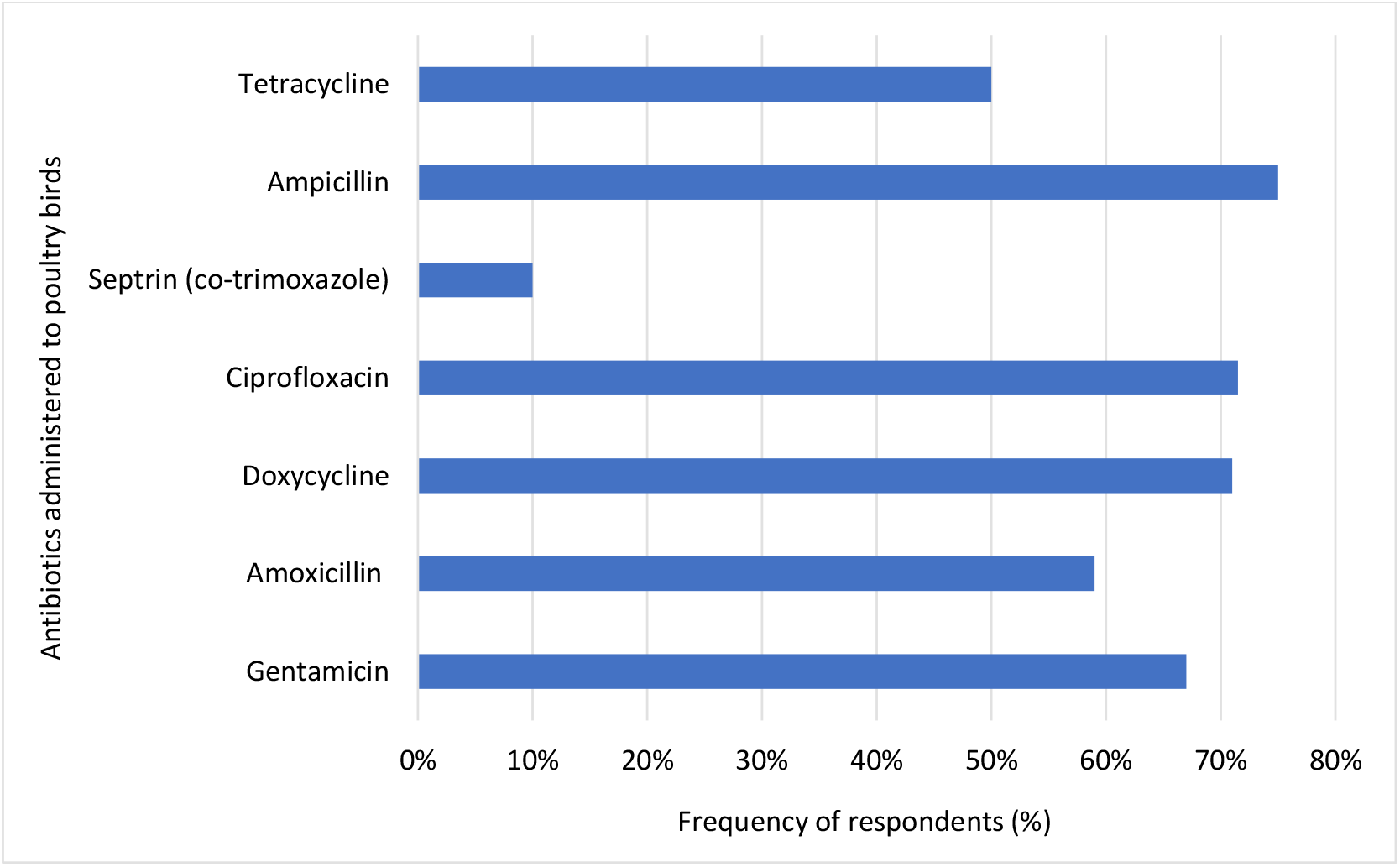
Commonly used antibiotics in poultry farms in Enugu, Nigeria.

**Figure 3.**
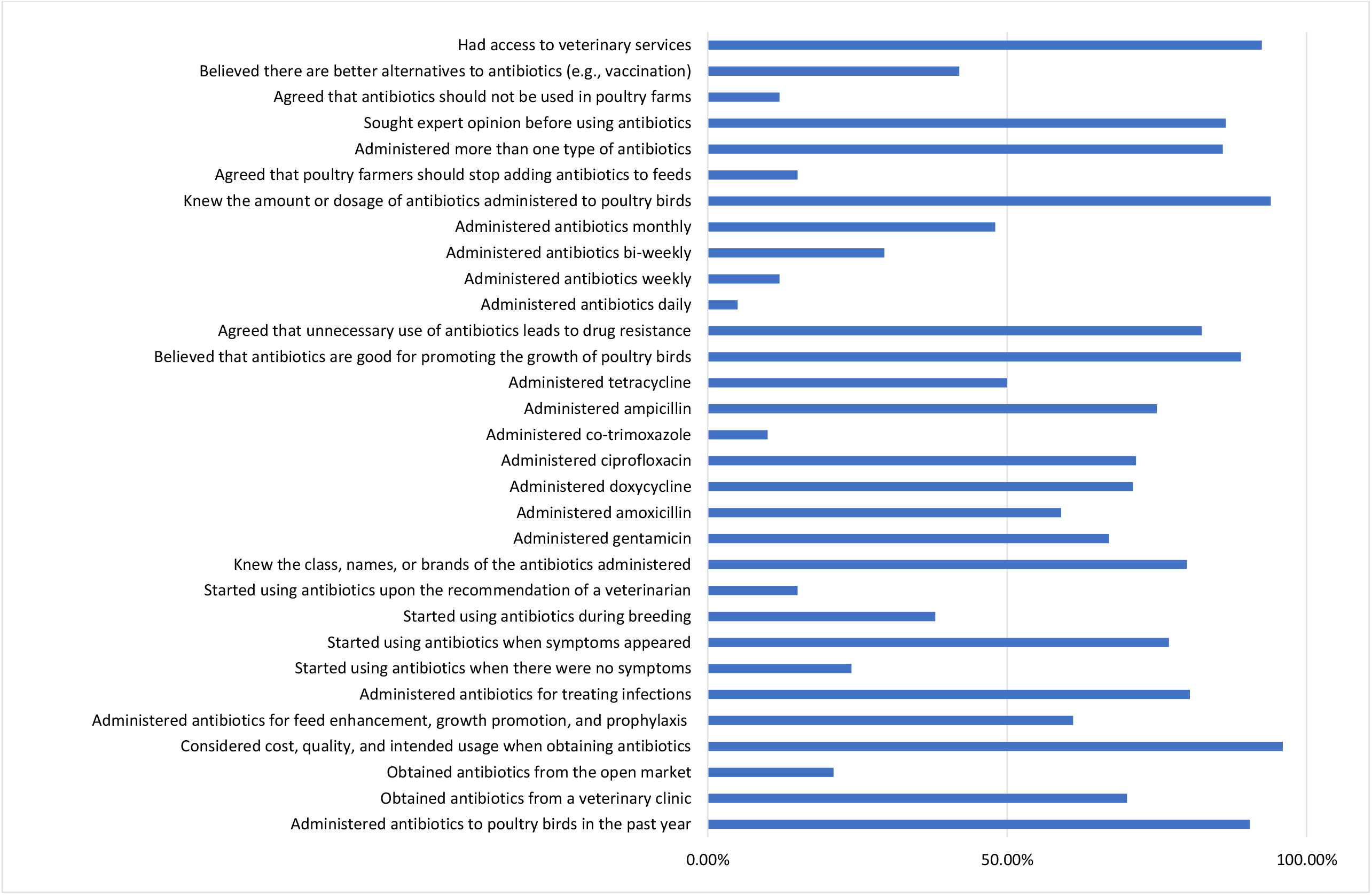
Distribution of insights into the KAP on antibiotics use by farmers in Enugu, Nigeria.

The insights into the KAP of poultry farmers’ regarding antibiotic use is presented in Figure 2. In terms of knowledge, 90.5% of respondents reported using antibiotics within the past year, primarily sourcing them from veterinary clinics (70%), with additional sources including pharmacies (23%) and open markets (21%). Though only 42% considered alternatives like vaccination viable, only 50.5% expressed uncertainty.

When asked if antibiotic application should be avoided in poultry farms 12% of the respondents agreed that antibiotics should be avoided on farms, with 74.5% disagreeing. A significant proportion of the respondents (86.5%) reported consulting experts like veterinarians before administering antibiotics to their birds. Nearly all respondents (96%) considered factors such as cost, quality, and intended use before purchasing any antibiotic. Surprisingly, 86% of the respondents used multiple antibiotics on their poultry birds, with 92.5% having access to veterinary services (Figure 2; Table S2).

### KAP of poultry farmers regarding AMR

The response of the poultry farmers on matters regarding their KAP on AMR is presented in Figure 3 and highlights key trends and gaps. When asked about awareness on AMR, a considerable portion of respondents (65%) were unaware of AMR, with only 33% indicating awareness. When asked about the role of antibiotics in promoting AMR, 42.5% of the respondents believed that antibiotic use in poultry farming contributes to resistance, while 22.5% disagreed, and 25% expressed uncertainty, reflecting an overall limited understanding (Figure 4; Table S3).

**Figure 4.**
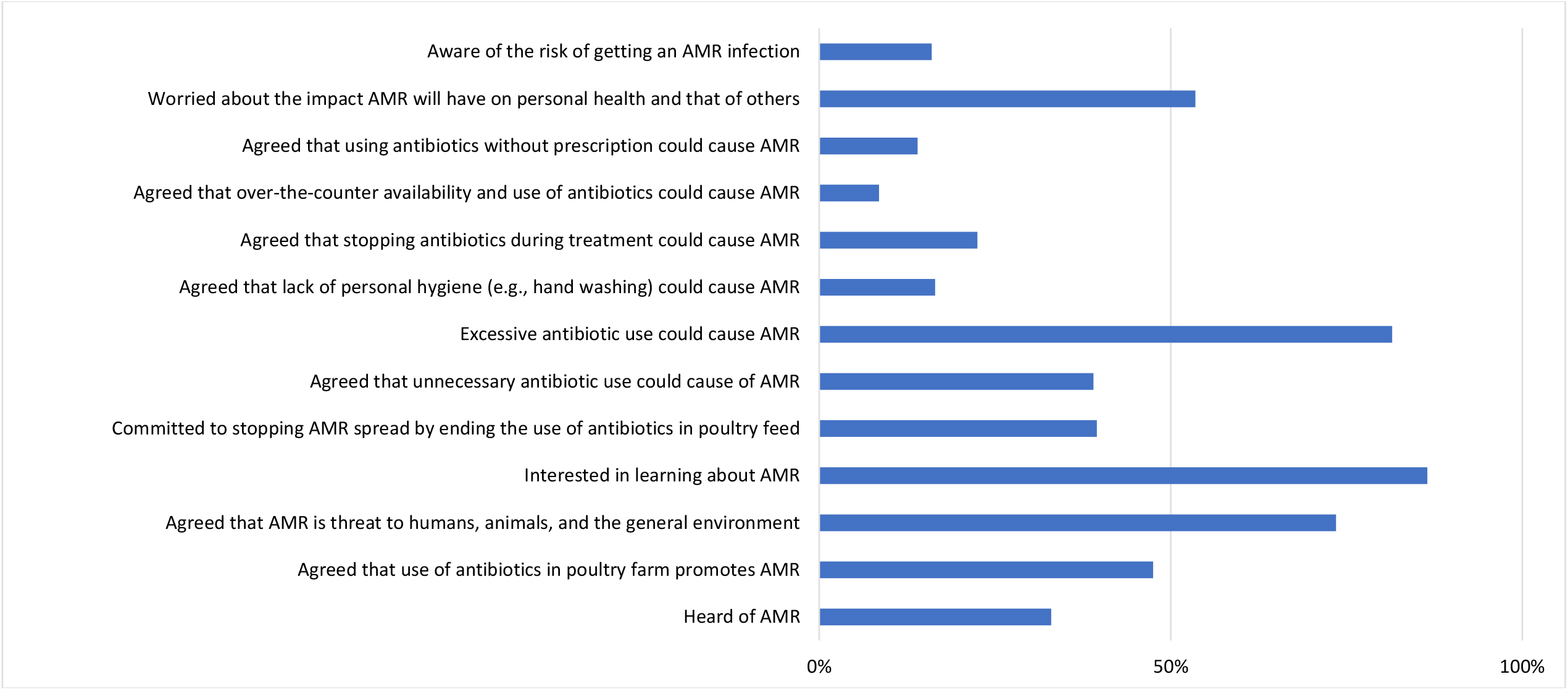
Distribution of insights into KAP on AMR by farmers in Enugu, Nigeria.

When asked if AMR is a threat to humans, animals and the environment, 55.5% of the respondents perceived AMR as a significant threat to human, animal, and environmental health. 22% of respondents are unaware with only 18% of the respondents strongly agreeing. In terms of attitude, 86.5% of farmers expressed interest in further education on AMR, yet only 39.5% were committed to ceasing antibiotic use in poultry feed, with 21% unwilling and 38% uncertain about making such changes (Figure 4; Table S3).

Regarding practice, 81.5% of the respondents identified excessive antibiotic use as a major contributing factor to AMR. This was followed by unnecessary use (39%) and treatment interruption (22.5%). Although 53.5% of respondents expressed concern about AMR’s health impacts, only 16% recognized a personal risk of acquiring an AMR infection, with 59.5% uncertain of their susceptibility (Figure 4; Table S3).

Regarding the relationship between poultry farmers’ knowledge of AMR and their socio-demographic factors - gender, age, marital status, occupation, education, and internet usage, there were no statistically significant risk factors influencing poor AMR awareness level for gender among the respondents. AMR knowledge levels are similar for males (51.14%) and females (42.86%), with no significant difference (p=0.410), indicating gender does not significantly affect AMR awareness.

However, there is significant difference in the knowledge level of respondents in the different age groups (p≤0.05). On the other hand, there is no significant difference in the knowledge level of respondents with different marital statuses (p>0.05). Regarding the relationship between respondent’s knowledge on AMR and other socio-demographic variables, there is significant difference in the knowledge level of respondents on AMR in terms of their occupational status, education and internet usage (p≤0.05) (Table S4).

Meanwhile, the younger respondents, particularly those aged 21-30, show the highest AMR knowledge (53.70%) compared to older groups, with a significant difference (p=0.005). Thus, age influences AMR awareness, with younger farmers likely benefiting from better access to recent information (Table S4).

On the other hand, single farmers have higher AMR knowledge (59.09%) than married farmers (40.15%), but this difference is not significant (p = 0.082), suggesting marital status alone is not a major factor in AMR awareness among the respondents. In this study, respondent’s knowledge on AMR varies significantly by occupation (p=0.006); employed respondents (58.11%) and students exhibit higher awareness compared to housewives (18.75%), likely due to greater exposure to information in work and educational settings (Table S4).

AMR knowledge increases with education level, from 0% for those with primary education to 87.5% for those with a doctorate (p=0.020). Higher education likely provides better access to information and critical thinking skills, influencing awareness on the subject matter. Daily or frequent internet users have significantly higher AMR knowledge (52.78% and 52.63%) compared to those with rare or no access (23.81% and 7.69%), with a significant difference (p=0.014), suggesting regular internet access is a key factor in AMR awareness (Table S4).

## DISCUSSION

AMR is increasing globally and poses a great health challenge if not halted. AMR in poultry farming represents a significant public health concern in Nigeria, where a lack of standardized guidelines for antibiotic use contributes to inconsistent practices across farms. Antibiotics are often readily accessible over the counter (OTC) and without prescription in the large part, leading to varied and unregulated usage patterns that can exacerbate AMR risks. This study addresses a critical gap by assessing the KAP of poultry farmers in Enugu, Nigeria, regarding antibiotic use and AMR.

A recent report highlights that AMR affects countries worldwide, irrespective of income levels, but is disproportionately exacerbated by poverty and inequality, with low- and middle-income countries (LMICs) bearing the greatest burden [16]. Inadequate training and education, combined with the misuse of antibiotics in poultry farming, are significant risk factors that further intensify the AMR crisis in LMICs, including Nigeria.

As previously reported, the incessant and unjustified application of antibiotics in animal production including poultry has dare consequences for human and environmental health especially as these antibiotics are closely related or the same as those used in human medicine [3,17]. In our study, there is a high level of antibiotic use among the poultry farmers/respondents. A significant majority (90.5%) of poultry farmers in this study reported administering antibiotics to their birds, with frequent administration patterns comprising monthly, bi-weekly, or weekly. Surprisingly, the poultry farmers relied on these antibiotics to treat infection rather than for growth. But on the other hand, a significant proportion of the respondents (89%) confirmed administering antibiotics to promote growth in poultry birds. This result corroborates with other studies conducted in India and Pakistan as it reveals that over 50% of antibiotic consumption in poultry farms were without prescription and applied to enhance growth in poultry birds [18, 19].

Still, majority of the farmers believe that antibiotics are beneficial for growth promotion (89%), and this indicates a widespread misunderstanding of appropriate antibiotic use in these milieus. Ampicillin (75%), followed by ciprofloxacin (71.5%), and doxycycline (71%) was identified as the most commonly used antibiotic in our study, raising concerns due to its potential link to beta-lactamase selection amid rising incidence of carbapenem resistance in Nigeria [1, 20]. These antibiotics were reported as the most common antibiotics applied in commercial poultry laying hens in Southwest Nigeria and Pakistan [19, 21].

Alhaji and Isola [22] also opined that antibiotics belonging to the beta-lactam, fluoroquinolone, aminoglycoside, sulfonamide, and tetracycline classes were amongst the widely used in food-animal production in Northcentral Nigeria. While the reasons for choosing these classes of antibiotics are still not clear, the OTC availability of antibiotics and previous common usage by farmers may be contributing factors driving AMR in poultry milieus. This continual irrational use of these classes of antibiotics in poultry farms increases the likelihood of developing AMR as bacteria are consistently exposed to these drugs, which can lead to the selection of resistant strains and genes since these antibiotics have similarities with those used in clinical medicine.

The most common sources of antibiotics identified in this study were veterinary clinics (70%), followed by pharmacies (23%), the open market (21%) and patent chemist shops (14.5%). While a significant proportion of respondents (86.5%) reported seeking expert opinions and services before deciding to use antibiotics, the availability of antibiotics OTC from these sources - often without a prescription -could exacerbate the AMR crisis in the studied region. In a related study conducted in Ethiopia, the primary sources of antibiotics were veterinary pharmacies (100%), veterinary clinics (51%), human pharmacies (26.8%), and open markets (16.2%) [23]. This trend was also observed in North-Central Nigeria, Southwest Nigeria, Burkina Faso, and Pakistan where similar sources of antibiotics for poultry farming have been reported [14, 19, 21, 24].

To ensure good practices, adequate knowledge of AMR among poultry farmers is crucial to prevent the irrational use of antibiotics - a major contributor to AMR prevalence. The level of AMR knowledge observed in this study is disturbing, with approximately 65% of respondents being unaware of AMR. This highlights a significant knowledge gap. Only 33% of respondents reported having heard of AMR. This aligns with findings from Reyher et al. [25], where 76.9% of participants reported a lack of AMR awareness. Similarly, a study in Ouagadougou, Burkina Faso, found that more than 80% of poultry farmers lacked knowledge of AMR [24], a finding comparable to ours (65%). This lack of awareness likely contributes to the continued use of antibiotics without sufficient consideration of their environmental impact, including the promotion of AMR. Overall, these findings suggest a basic awareness of AMR’s threat, but knowledge gaps, limited commitment to behavioral change, and confusion about risk factors and causative practices remain. These barriers may hinder efforts to mitigate AMR in poultry farming in Nigeria.

Even among those aware of AMR, confusion persists regarding its causes and personal risk factors. While 47.5% recognize that antibiotic use in poultry farming contributes to AMR, a notable portion (26%) does not see the connection, and 25% remain uncertain. This lack of clarity may result in practices that unintentionally promote the spread of AMR. Our study found that awareness of AMR is influenced by factors such as age, education, occupation, and internet access, with younger, more educated farmers who use the internet more likely to be aware. This lack of awareness, combined with high antibiotic use, contributes to the spread of AMR from poultry to humans and the environment, emphasizing the need for targeted education and interventions.

Among male respondents, 51.14% demonstrated knowledge of AMR, compared to 42.86% of female respondents; however, the p-value of 0.410 suggests no statistically significant difference. Similarly, 59.09% of single respondents had greater AMR knowledge than married respondents (40.15%), with a p-value of 0.082, indicating no significant effect of marital status. These results align with previous studies, highlighting that AMR knowledge is more strongly associated with access to information and education rather than gender or marital status.

AMR knowledge also varied by age group, with the highest level (53.70%) observed in individuals aged 21-30 and the lowest (30.43%) in those over 60 (p = 0.005). Occupational status influenced AMR knowledge, with employed respondents demonstrating the highest knowledge levels (58.11%) and housewives the lowest (18.75%) (p = 0.006). Education level was strongly correlated with AMR knowledge, as respondents with a master’s (83.33%) or doctorate (87.5%) had the highest knowledge, while those with only a primary education showed the lowest levels (0%) (p = 0.020). In conclusion, demographic factors such as age, employment status, educational attainment, and internet usage significantly affect AMR knowledge. Younger, employed, more educated individuals, and frequent internet users exhibited higher awareness, likely due to better access to information and educational resources.

Overall, the data from our study underscore the importance of adopting non-pharmaceutical approaches to prevent infection in poultry settings. These include implementing strong biosecurity policies, maintaining proper personal and environmental hygiene, ensuring the correct separation of birds, using clean water and feeding troughs, and reducing antibiotic use. Such practices are vital in mitigating AMR, as poultry farming is a significant source of AMR due to the widespread use of antibiotics aimed at enhancing production through modern practices [18]. These findings highlight the need for targeted AMR-related educational programs, particularly for poultry farmers. Tailoring outreach strategies and providing offline resources can enhance AMR awareness, empowering individuals to engage in responsible antibiotic practices, which is crucial for mitigating the spread of AMR in Nigeria, where antibiotics are readily available OTC.

Although the study focused on investigating antibiotic use and AMR in poultry farms within a single state (Enugu State) in Nigeria, providing valuable insights into local sub-communities there, its findings offer a broader descriptive overview of KAP related to antibiotic use and AMR on poultry farms across Nigeria. This research underscores the risks posed by farmers’ inadequate knowledge of antibiotics and AMR, highlighting potential hazards to human health, animal welfare, and the environment. It emphasizes the urgent need for strict antibiotic policies in poultry farming and the development of targeted education, training, and awareness programs for all stakeholders in the poultry industry. These measures are essential for mitigating the emergence and spread of AMR in local communities across Nigeria.

## CONCLUSION

AMR in poultry farming poses a critical public health challenge in Nigeria, driven by widespread misuse of antibiotics, limited awareness among farmers, and unregulated over-the-counter availability of antibiotics. Our study reveals that 90.5% of poultry farmers frequently use antibiotics, often without prescriptions, while only 33% are aware of AMR and its risks, highlighting significant knowledge gaps. The most striking finding in our study is the heavy reliance on antibiotics by poultry farmers, primarily for treating infections and, to a significant extent, for promoting growth. This reliance persists despite limited awareness of AMR. Ampicillin was identified as the most commonly used antibiotic, raising concerns due to its potential contribution to beta-lactamase selection, particularly in the context of Nigeria’s growing carbapenem resistance issues. These findings underscore the urgent need for targeted education, stricter regulation of antibiotic use in poultry farming, and the adoption of non-pharmaceutical practices to mitigate AMR spread and safeguard human, animal, and environmental health.

## Acknowledgment

We acknowledge the management of Enugu State University of Science and Technology (ESUT), Agbani, Nigeria and the Administrative/Technical staff from ACEGID, Redeemer’s University, Ede, Osun State, Nigeria for all administrative support towards the successful conduct and completion of this study.

## Supplementary Material

The following Tables – S1: Relationship between respondents’ knowledge about antibiotic use and their socio-demographic data; Table S2: Knowledge, Attitude and Practice on Antibiotics Use; Table S3: Knowledge, Attitude and Practice on Antimicrobial Resistance (AMR); and Table S4: Relationship between respondents’ knowledge about AMR and their socio-demographic data – are available in in the supplementary material.

## Authors contribution

Conceptualization: CPE

Methodology: CPE, CWA, PME

Formal analysis: EAN, ENO, MUA, PME

Data analysis: CPE, EAN, PME Supervision: CPE, MUA, PME

Writing, review and editing: CPE, EAN, CWA, ENO, MUA, PME

