## Supplementary Tables 1-4 for "Investigating the factors influencing antibiotic use practices and their association with antimicrobial resistance awareness among poultry farmers in Enugu State, Nigeria"

### Table of content


| | | |
|---|---|---|
| **Table S1:** | Relationship between respondents' knowledge about antibiotic use and their socio-demographic data | **3** |
| **Table S2:** | Knowledge, Attitude and Practice on Antibiotics Use | **4** |
| **Table S3:** | Knowledge, Attitude and Practice on Antimicrobial Resistance (AMR) | **6** |
| **Table S4:** | Relationship between respondents' knowledge about AMR and their socio-demographic data | **7** |

**Table S1: Relationship between respondents' knowledge about antibiotic use and their socio-demographic data**

| **Demographic variable** | | Knowledge level | | | | **Total** | | **p-value** (calculated using 2-way ANOVA) | Remark |
|---|---|---|---|---|---|---|---|---|---|
| | | **No** | | **Yes** | | | | | |
| | | n | % | n | % | n | % | | |
| **Gender** | Male | 6 | 6.82 | 82 | 93.18 | 88 | 100 | 0.570 | p>0.05 (There is no significant difference in the knowledge level of the male and female respondents). |
| | Female | 3 | 2.68 | 109 | 97.32 | 112 | 100 | | |
| **Age** | <20 years | 0 | 0 | 8 | 100 | 8 | 100 | 0.392 | p>0.05 (There is no significant difference in the knowledge level of respondents in the different age groups). |
| | 21 – 30 years | 3 | 5.56 | 51 | 94.44 | 54 | 100 | | |
| | 31 – 40 years | 1 | 2 | 49 | 98 | 50 | 100 | | |
| | 41 – 50 years | 4 | 9.09 | 40 | 90.91 | 44 | 100 | | |
| | 51-60 years | 0 | 0 | 21 | 100 | 21 | 100 | | |
| | >60 years | 1 | 4.35 | 22 | 95.65 | 23 | 100 | | |
| **Marital Status** | Single | 6 | 9.09 | 60 | 90.91 | 66 | 100 | 0.487 | p>0.05 (There is no significant difference in the knowledge level of respondents with different marital statuses). |
| | Married | 2 | 1.52 | 130 | 98.48 | 132 | 100 | | |
| | Divorced | 0 | 0 | 1 | 100 | 1 | 100 | | |
| **Occupational Status** | Employed | 2 | 2.70 | 72 | 97.30 | 74 | 100 | 0.448 | P>0.05 (There is no significant difference in the knowledge level of respondents with different occupational statuses). |
| | Self Employed | 1 | 2.38 | 41 | 97.62 | 42 | 100 | | |
| | Housewife | 0 | 0 | 16 | 100 | 16 | 100 | | |
| | Househusband | 0 | 0 | 1 | 100 | 1 | 100 | | |
| | Student | 3 | 8.33 | 33 | 91.67 | 36 | 100 | | |
| | Retired | 1 | 6.25 | 15 | 93.75 | 16 | 100 | | |
| | Others | 1 | 7.14 | 13 | 92.86 | 14 | 100 | | |
| **Highest Academic Qualifications** | Primary school | 1 | 20 | 4 | 80 | 5 | 100 | 0.457 | P>0.05 (There is no significant difference in the knowledge level of respondents with different academic qualifications). |
| | Secondary school | 1 | 1.61 | 61 | 98.39 | 62 | 100 | | |
| | University | 3 | 3.95 | 73 | 96.05 | 76 | 100 | | |
| | Diploma | 2 | 16.67 | 10 | 83.33 | 12 | 100 | | |
| | OND/HND | 0 | 0 | 11 | 100 | 11 | 100 | | |
| | Masters | 1 | 5.56 | 17 | 94.44 | 18 | 100 | | |
| | Doctorate (PhD) | 0 | 0 | 8 | 100 | 8 | 100 | | |
| **Internet Usage** | Everyday | 4 | 3.70 | 104 | 96.30 | 108 | 100 | 0.467 | P>0.05 (There is no significant difference in the knowledge level of respondents with varying degrees of internet use). |
| | Often | 1 | 1.75 | 56 | 98.25 | 57 | 100 | | |
| | Rarely | 2 | 9.52 | 19 | 90.48 | 21 | 100 | | |
| | Never | 1 | 7.69 | 12 | 92.31 | 13 | 100 | | |

**Table S2: Knowledge, Attitude and Practice on Antibiotics Use**

| **Variable** | **Response** | **n** | **%** |
|---|---|---|---|
| Did you administer antibiotics to your birds within the past year? | Yes | 181 | 90.5 |
| | No | 19 | 9.5 |
| Where did you obtain the antibiotics that you have administered within the past year? | Patent chemist shop | 29 | 14.5 |
| | Pharmacy | 46 | 23 |
| | Open market | 42 | 21 |
| | Drug vendors | 12 | 6 |
| | Hospital | 37 | 18.5 |
| | Veterinary clinic | 140 | 70 |
| Do you consider cost, quality and the intended usage when buying antibiotics for your birds? | Yes | 192 | 96 |
| | No | 4 | 2 |
| | No idea | 2 | 1 |
| | No response | 2 | 1 |
| What was your reason for administering antibiotics to your birds? | For growth promotion | 25 | 12.5 |
| | For prophylaxis | 87 | 43.5 |
| | For treating infection | 161 | 80.5 |
| | For feed enhancement | 10 | 5 |
| | Other | 3 | 1.5 |
| When do you start the use of antibiotics? | When there are no symptoms of infection | 48 | 24 |
| | When there are symptoms of infection | 154 | 77 |
| | During breeding | 76 | 38 |
| | Upon the recommendation of a veterinarian | 30 | 15 |
| | At all times | 5 | 2.5 |
| | When I feel like | 4 | 2 |
| Did you know the class, names or brands of the antibiotics you administered to your birds | Yes | 160 | 80 |
| | No | 40 | 20 |
| List of the antibiotics class or names | Gentamicin | 134 | 67 |
| | Amoxicillin | 118 | 59 |
| | Doxycycline | 142 | 71 |
| | Ciprofloxacin | 143 | 71.5 |
| | Septrin (co-trimoxazole) | 20 | 10 |
| | Ampicillin | 150 | 75 |
| | Tetracycline | 100 | 50 |
| | No response | 41 | 20.5 |
| Antibiotics are good for promoting the growth of poultry birds | Correct | 178 | 89 |
| | Wrong | 5 | 2.5 |
| | I don’t know | 14 | 7 |
| | No response | 3 | 1.5 |
| Unnecessary use of antibiotics leads to that drug losing its effectiveness in future | Yes | 165 | 82.5 |
| | No | 7 | 3.5 |
| | I don’t know | 24 | 12 |
| | No response | 4 | 2 |
| How frequent do you administer antibiotics? | Daily | 10 | 5 |
| | Weekly | 24 | 12 |
| | Bi-weekly | 59 | 29.5 |
| | Monthly | 96 | 48 |
| | Yearly | 4 | 2 |
| | No response | 7 | 3.5 |
| Do you know the amount or dosage of antibiotics administered? | Yes | 188 | 94 |
| | No | 5 | 2.5 |
| | No response | 7 | 3.5 |
| Poultry farmers should stop introducing antibiotics in feeds? | Agree | 30 | 15 |
| | Disagree | 144 | 72 |
| | I don’t know | 21 | 10.5 |
| | No response | 5 | 2.5 |

| | | | |
|---|---|---|---|
| Do you give your poultry birds more than one type of antibiotics? | Yes | 172 | 86 |
| | No | 16 | 8 |
| | I don't know | 5 | 2.5 |
| | No response | 7 | 3.5 |
| Do you seek for the services/opinion of an expert before deciding to use antibiotics? | Yes | 173 | 86.5 |
| | No | 19 | 9.5 |
| | No response | 8 | 4 |
| Do you agree that antibiotics should not be used in poultry farms? | Yes | 24 | 12 |
| | No | 149 | 74.5 |
| | I don't know | 22 | 11 |
| | No response | 5 | 2.5 |
| Do you agree that there are better alternatives (e.g., vaccination) to the use of antibiotics? | Yes | 84 | 42 |
| | No | 10 | 5 |
| | I don't know | 101 | 50.5 |
| | No response | 5 | 2.5 |
| Do you have access to veterinary services? | Yes | 185 | 92.5 |
| | No | 10 | 5 |
| | No response | 5 | 2.5 |

**Table S3: Knowledge, Attitude and Practice on Antimicrobial Resistance (AMR)**

| Variable | Response | n | % |
|---|---|---|---|
| Have you heard of antimicrobial resistance (AMR)? | Yes | 66 | 33 |
| | No | 130 | 65 |
| | No response | 4 | 2 |
| The use of antibiotics in poultry farm promotes AMR | Strongly Agree | 10 | 5 |
| | Agree | 85 | 42.5 |
| | Disagree | 45 | 22.5 |
| | Strongly disagree | 7 | 3.5 |
| | I don't know | 50 | 25 |
| | No response | 3 | 1.5 |
| AMR is threat to humans, animals and the general environment | Strongly Agree | 36 | 18 |
| | Agree | 111 | 55.5 |
| | Disagree | 4 | 2 |
| | Strongly disagree | 2 | 1 |
| | I don't know | 44 | 22 |
| | No response | 3 | 1.5 |
| How interested are you to learn about AMR? | Interested | 173 | 86.5 |
| | Not interested | 6 | 3 |
| | Undecided | 18 | 9 |
| | No response | 3 | 1.5 |
| Are you committed to stop AMR spread by ending the use of antibiotics in poultry feed? | Yes | 79 | 39.5 |
| | No | 42 | 21 |
| | I don't know | 76 | 38 |
| | No response | 3 | 1.5 |
| What do you think could cause AMR? Please select or tick all that you think applies | Unnecessary antibiotic use | 78 | 39 |
| | Excessive antibiotic use | 163 | 81.5 |
| | Lack of personal hygiene (e.g., hand washing) | 33 | 16.5 |
| | Stoppage of antibiotics during treatment | 45 | 22.5 |
| | Over the counter availability and use of antibiotics | 17 | 8.5 |
| | Using antibiotics without prescription | 28 | 14 |
| | Others | 16 | 8 |
| | I don't know | 21 | 10.5 |
| Are you worried about the impact AMR will have on your health and that of others? | Yes | 107 | 53.5 |
| | No | 46 | 23 |
| | I don't know | 44 | 22 |
| | No response | 3 | 1.5 |
| Are you at risk of getting an AMR infection? | Yes | 32 | 16 |
| | No | 46 | 23 |
| | I don't know | 119 | 59.5 |
| | No response | 3 | 1.5 |

**Table S4: Relationship between respondents' knowledge about AMR and their socio-demographic data**

| Demographic variable | | Knowledge level No n | No % | Yes n | Yes % | Total n | Total % | p-value (calculated using 2-way ANOVA) | Remark |
|---|---|---|---|---|---|---|---|---|---|
| **Gender** | Male | 43 | 48.86 | 45 | 51.14 | 88 | 100 | 0.410 | p>0.05 (There is no significant difference in the knowledge level of the male and female respondents). |
| | Female | 64 | 57.14 | 48 | 42.86 | 112 | 100 | | |
| **Age** | <20 years | 4 | 50 | 4 | 50 | 8 | 100 | 0.005 | P≤0.05 (There is significant difference in the knowledge level of respondents in the different age groups). |
| | 21 – 30 years | 25 | 46.30 | 29 | 53.70 | 54 | 100 | | |
| | 31 – 40 years | 28 | 56 | 22 | 44 | 50 | 100 | | |
| | 41 – 50 years | 24 | 54.55 | 20 | 45.45 | 44 | 100 | | |
| | 51-60 years | 10 | 47.62 | 11 | 52.38 | 21 | 100 | | |
| | >60 years | 16 | 69.57 | 7 | 30.43 | 23 | 100 | | |
| **Marital Status** | Single | 27 | 40.91 | 39 | 59.09 | 66 | 100 | 0.082 | p>0.05 (There is no significant difference in the knowledge level of respondents with different marital statuses). |
| | Married | 79 | 59.85 | 53 | 40.15 | 132 | 100 | | |
| | Divorced | 0 | 0 | 1 | 100 | 1 | 100 | | |
| **Occupational Status** | Employed | 31 | 41.89 | 43 | 58.11 | 74 | 100 | 0.006 | P≤0.05 (There is significant difference in the knowledge level of respondents with different occupational statuses). |
| | Self Employed | 23 | 54.76 | 19 | 45.24 | 42 | 100 | | |
| | Housewife | 13 | 81.25 | 3 | 18.75 | 16 | 100 | | |
| | Househusband | 0 | 0 | 1 | 100 | 1 | 100 | | |
| | Student | 17 | 47.22 | 19 | 52.78 | 36 | 100 | | |
| | Retired | 11 | 68.75 | 5 | 31.25 | 16 | 100 | | |
| | Others | 11 | 78.57 | 3 | 21.43 | 14 | 100 | | |
| **Highest Academic Qualification** | Primary school | 5 | 100 | 0 | 0 | 5 | 100 | 0.020 | P≤0.05 (There is significant difference in the knowledge level of respondents with different academic qualifications). |
| | Secondary school | 43 | 69.35 | 19 | 30.65 | 62 | 100 | | |
| | University | 37 | 48.68 | 39 | 51.32 | 76 | 100 | | |
| | Diploma | 5 | 41.67 | 7 | 58.33 | 12 | 100 | | |
| | OND/HND | 7 | 63.64 | 4 | 36.36 | 11 | 100 | | |
| | Masters | 3 | 16.67 | 15 | 83.33 | 18 | 100 | | |
| | Doctorate (PhD) | 1 | 12.5 | 7 | 87.5 | 8 | 100 | | |
| **Internet Usage** | Everyday | 51 | 47.22 | 57 | 52.78 | 108 | 100 | 0.014 | P≤0.05 (There is significant difference in the knowledge level of respondents with varying degrees of internet use). |
| | Often | 27 | 47.37 | 30 | 52.63 | 57 | 100 | | |
| | Rarely | 16 | 76.19 | 5 | 23.81 | 21 | 100 | | |
| | Never | 12 | 92.31 | 1 | 7.69 | 13 | 100 | | |
